# Counting with DNA in metabarcoding studies: how should we convert sequence reads to dietary data?

**DOI:** 10.1101/303461

**Authors:** Bruce E. Deagle, Austen C. Thomas, Julie C. McInnes, Laurence J. Clarket, Eero J. Vesterinen, Elizabeth L. Clare, Tyler R. Kartzinel, J. Paige Eveson

## Abstract

Advances in DNA sequencing technology have revolutionised the field of molecular analysis of trophic interactions and it is now possible to recover counts of food DNA barcode sequences from a wide range of dietary samples. But what do these counts mean? To obtain an accurate estimate of a consumer’s diet should we work strictly with datasets summarising the frequency of occurrence of different food taxa, or is it possible to use the relative number of sequences? Both approaches are applied in the dietary metabarcoding literature, but occurrence data is often promoted as a more conservative and reliable option due to taxa-specific biases in recovery of sequences. Here, we point out that diet summaries based on occurrence data overestimate the importance of food consumed in small quantities (potentially including low-level contaminants) and are sensitive to the count threshold used to define an occurrence. Our simulations indicate that even with recovery biases incorporated, using relative read abundance (RRA) information can provide a more accurate view of population-level diet in many scenarios. The ideas presented here highlight the need to consider all sources of bias and to justify the methods used to interpret count data in dietary metabarcoding studies. We encourage researchers to continue to addressing methodological challenges, and acknowledge unanswered questions to help spur future investigations in this rapidly developing area of research.

## 1. Introduction

Many recent studies documenting trophic interactions make use of metabarcoding, an approach which combines high-throughput sequencing (HTS) with DNA barcoding to characterise organisms in complex mixtures (Nielsen *et al.* 2017). When HTS first became available the potential applications in diet studies were clear and the methods were quickly embraced by the community (Deagle *et al.* 2009; Valentini *et al.* 2009). In a comprehensive review of DNA-based diet analysis by King *et al.* (2008) the possibility of using HTS was only briefly mentioned as a ‘Future Direction’, and just four years later another review paper focussed entirely on this approach (Pompanon *et al.* 2012). While many underlying technical and biological details vary between dietary metabarcoding studies, the general workflow is now well defined. It involves extraction of DNA from faecal samples or stomach contents, PCR amplification of DNA barcode markers from food taxa of interest, and then DNA sequencing for taxonomic classification of the recovered sequences. The workflow has been applied to determine diet in a range of animals, from invertebrates to large mammalian herbivores and carnivores (representative studies summarised in Table 1).

**Table 1.**
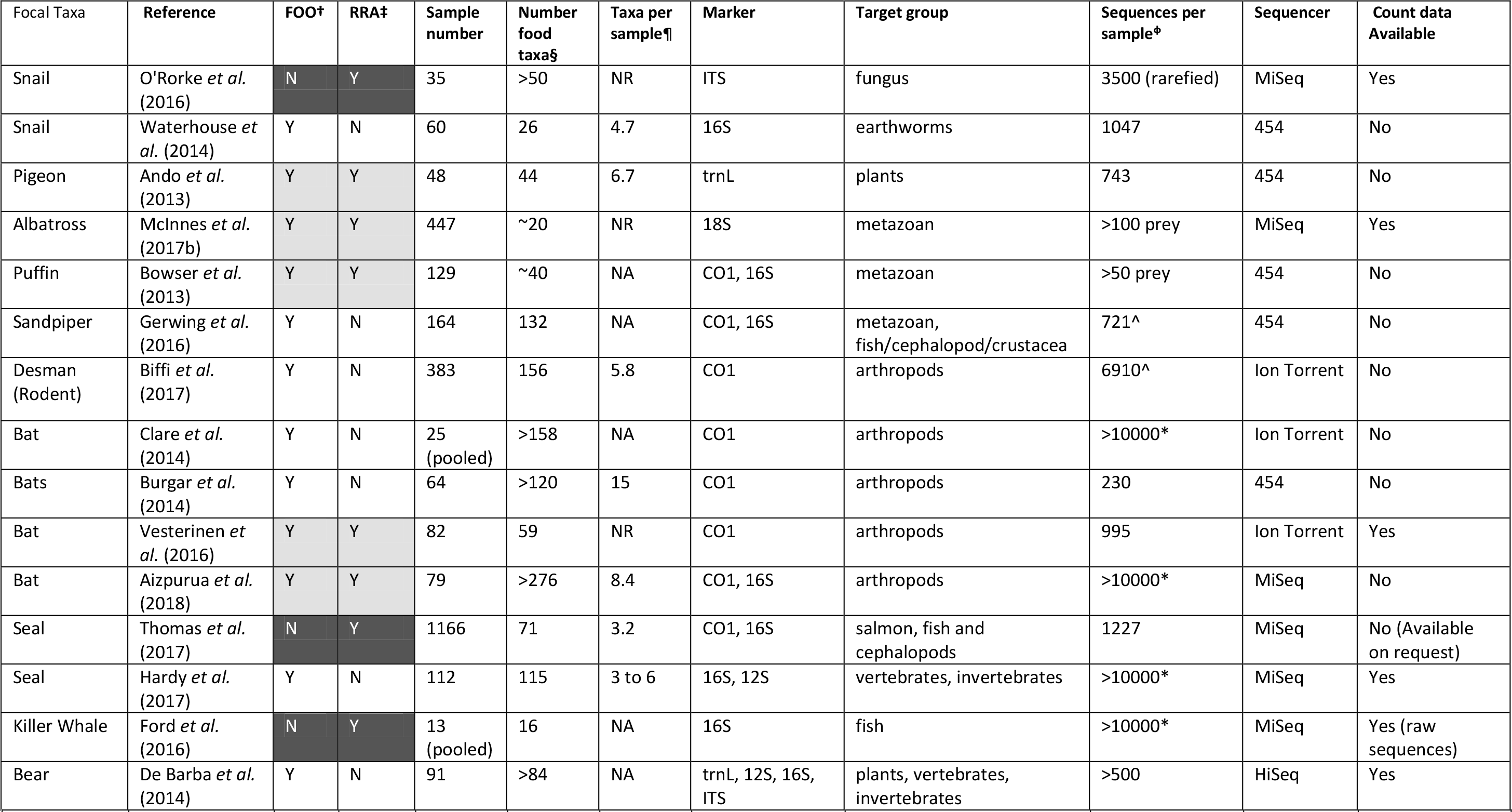
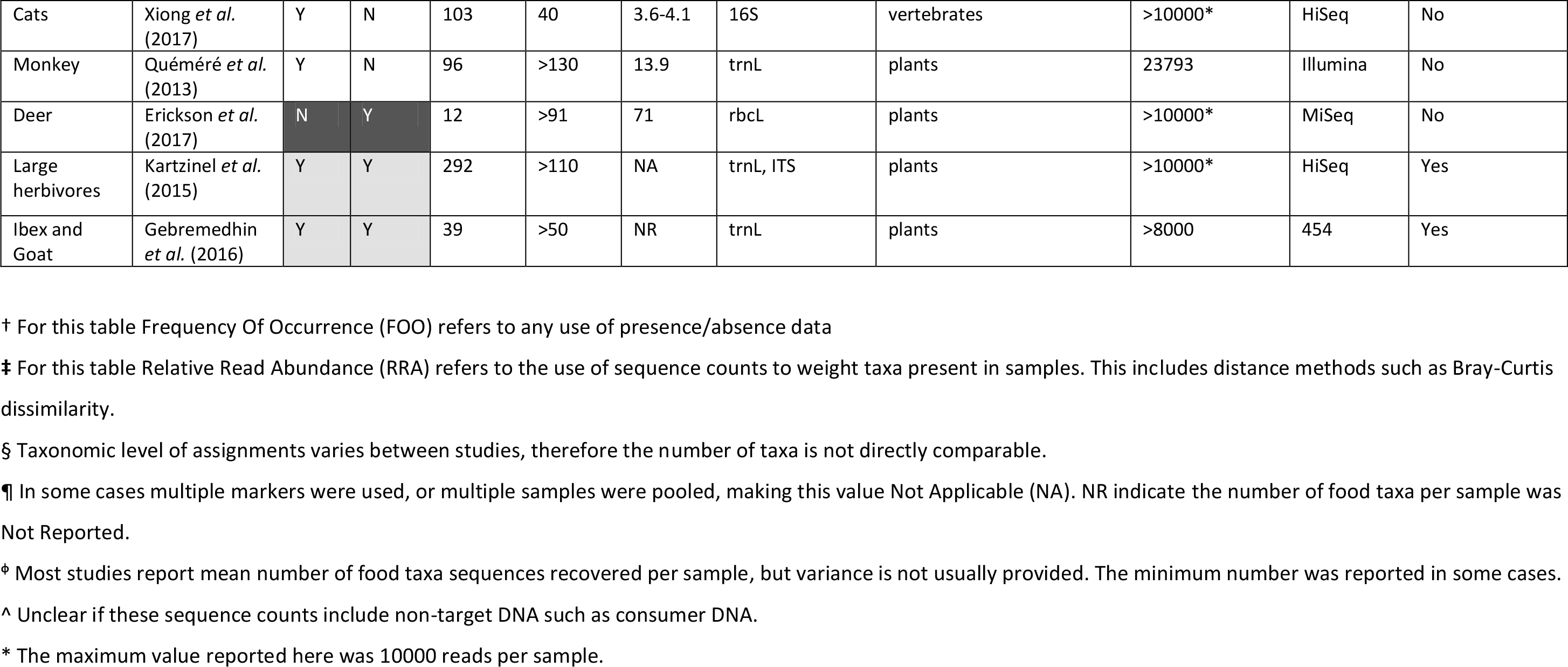
Use of sequence counts in 20 metabarcoding diet studies carried out using faecal DNA collected from a range of different species. Representative studies across a range of focal taxa carried out by different research groups are shown rather than trying to summarise all dietary metabarcoding studies.

The rapid adoption of HTS to characterise complex mixtures of DNA is not unique to dietary studies; over the last ten years the technology has produced a wealth of new genetic data providing insight into almost all areas of biology (Goodwin *et al.* 2016). One feature of HTS is that it provides counts of DNA sequences in each sample and therefore it has the potential not only to provide a qualitative list, but also to quantify what DNA is present. The interpretation of sequence read counts as a numerical representation of sample composition is common in many HTS applications. For example, studies sequencing transcripts to determine differences in gene expression (Finotello & Di Camillo 2015), profiling microbe communities (Vandeputte *et al.* 2017) or measuring epigenetic variation (Schield *et al.* 2016) all rely on sequence read counts. However, decisions about how to interpret read counts is certainly not routine and the validity of interpretations is sometimes questioned even in fields where the practice is well established (e.g. Edgar 2017; Olova *et al.* 2017). These debates are constructive, and should motivate researchers to test the underlying assumptions and justify their interpretations, but can inadvertently give rise to the false impression that count data are always misleading.

The reality is that metabarcoding studies always use sequence counts to some extent. In dietary investigations, count data are used either to record the occurrence of food species within samples based on a threshold number of sequences (i.e. presence/absence of taxa), or to calculate the percentage of DNA belonging to each food species as a proxy for relative biomass consumed (i.e. relative abundance of taxa; Figure 1). The conversion of sequence counts to occurrence data is often considered a more conservative approach than using proportional data. In their introduction to the Molecular Ecology Special Issue on ‘Molecular Detection of Trophic Interactions’, Symondson & Harwood (2014) pointed out that authors of many metabarcoding papers “*now simply record numbers of predators testing positive for a target prey or plant species, providing a pragmatic and useful surrogate for truly quantitative information*”. This sentiment, that focusing only on occurrence data is a reliable and safe option, is now common in the literature and this step in the analysis pipeline is often uncritically applied as the default option. Using counts as an indication of biomass in the sample is more controversial. Indeed, the difficulties of obtaining an accurate biomass signature from sequence counts include both technical and biological biases that affect barcode marker recovery rates from different taxa (Amend *et al.* 2010; Deagle *et al.* 2009; Pompanon *et al.* 2012). Therefore in the best-case scenario sequence read counts can only provide a rough estimate of proportional abundance. Still, to accept the notion that relative sequence counts provide no meaningful information would mean that, within one sample, a few DNA sequences from one food taxon is equivalent to 10,000 sequences from another. Most molecular ecologists would interpret these disparate counts to mean that there are differences in template DNA abundance (as long as methods used to collect the data are reasonable) and that there is some biological basis for that difference. Ignoring this difference may inhibit ecological understanding.

**Figure 1:**
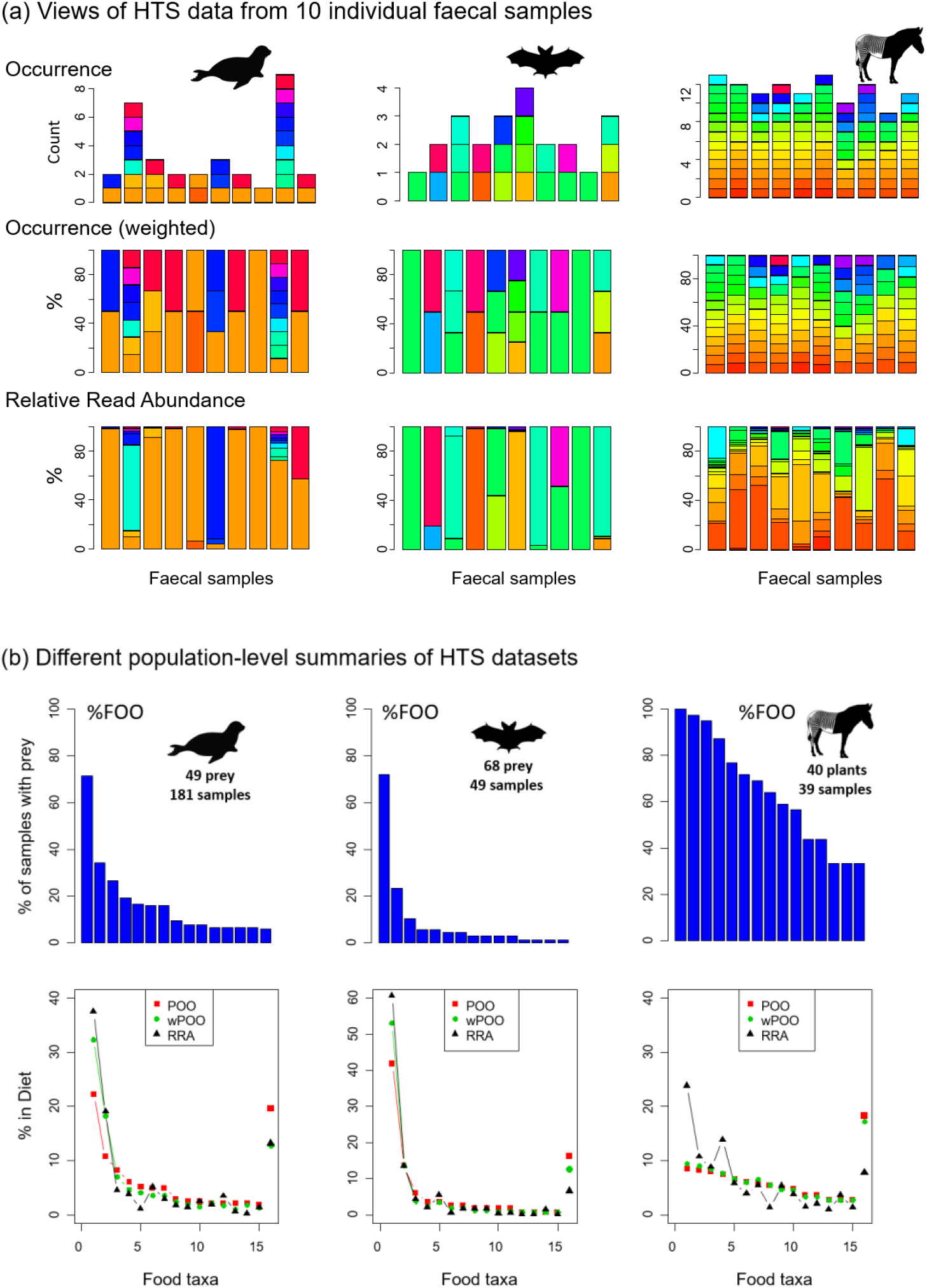
Information in faecal samples from dietary metabarcoding datasets of harbour seal (Thomas *et al.* 2017), an insectivorous bat (Vesterinen *et al.* 2016) and Grevy’s zebra (Kartzinel *et al.* 2015). (a) Individual-level data in 10 faecal samples viewed using different metrics. Colours represent different food taxa. (b) Population-level summaries of these datasets showing the top 15 food taxa (%FOO ranking); 1% threshold used for occurrence in POO and wPOO calculations. In the lower plots the sum contribution of remaining food taxa are plotted at end. In each example population data include only collections from one site and samples with >50 food taxa reads.

Here, we review the approaches taken to interpret sequence count data in dietary metabarcoding studies and consider their implications. We point out that converting sequence read counts to occurrence information can introduce strong biases and thus we suggest it is not always a “conservative” approach. We also assess the scale of biases in recovery of sequences from different food taxa in study systems where it has been examined. Using simulations we explore the impact of these biases on data summaries (both based on occurrence and read counts). In this light, we evaluate factors that impact data summaries in dietary metabarcoding and consider where using sequence count data as an indication of relative biomass within samples might be justified to provide a more nuanced picture of animal diet.

The issues we consider on how best to summarise dietary data have implications for all metabarcoding studies (Taberlet *et al.* 2018) and similar issues have been considered extensively in traditional diet studies (e.g. Barrett *et al.* 2007; Laake *et al.* 2002). In HTS-based diet studies the ideas are most relevant when the underlying objective is to estimate the true diet of a particular consumer (i.e. the relative biomass contributions of alternative diet species). This may not be a clearly stated goal, but is often implicit in outcomes of dietary metabarcoding studies. Approaches for summarising sequence counts may be of less concern in studies aiming to provide a list of taxa consumed by a particular species (niche breadth), a summary of trophic interactions in a food web, or an indicator of dietary differences between sites. Throughout the paper we will refer to the two general approaches of summarising sequence count data as ‘occurrence’ (i.e. presence/absence of taxa) and ‘relative read abundance’ (RRA; i.e. proportional summaries of counts). We focus mainly on dietary studies using DNA extracted from faecal material. The use of HTS to identify food in stomach contents is common in invertebrates, and also fish, but the material recovered is in various states of digestion and the sequence counts are less likely to contain a meaningful quantitative signal compared to the more consistent signal seen in faecal material (Deagle *et al.* 2013; Nakahara *et al.* 2015).

## 2. Current Practice

Non-dietary metabarcoding studies use a range of approaches to interpret sequence count data, and these vary depending on the targeted organisms. Recent papers published in Molecular Ecology on bacterial/archaeal communities all make use of RRA, although half of these studies also presented summaries based on taxon occurrences (Table S1). There is widespread acknowledgement of taxon-specific biases in recovery of the bacterial/archaeal barcode markers, but RRA is accepted as a flawed, but useful, measure of these diverse communities that cannot be easily characterized by other means (Forney *et al.* 2004; Ibarbalz *et al.* 2014). There is no clear consensus in metabarcoding of eukaryotic communities: RRA is sometimes used exclusively (often the case in studies of fungi), whereas metazoan studies use either occurrence data only or both metrics in tandem (recent examples listed in Table S1).

In dietary metabarcoding studies, it is common to only interpret sequence data after conversion to taxon occurrences (representative studies summarised in Table 1). This conversion is done in various ways. During initial processing of sequence reads, most researchers discard rare sequences to avoid incorporation of background sequencing errors (e.g. Quéméré *et al.* 2013). After this a summary table of remaining sequence reads in each sample is produced and sequences are assigned taxonomy (often with similar sequences being clustered). Then, when converting these read counts to occurrence data, a threshold number of reads is often required for each taxon to be tallied as an occurrence. Sequencing depth can vary considerably between samples, so in practice a threshold percentage of reads is often used (e.g. 1% of food sequences McInnes *et al.* 2017b), or sequencing depth can be rarefied to a common level (O’Rorke *et al.* 2016). These approaches normalize detection across samples, so that more sequences are required for an occurrence to be recorded in samples with higher read depths.

Once occurrences are recorded in individual samples, several metrics can be used to summarise the diet across samples. Those considered here are percent frequency of occurrence (%FOO), percent of occurrence (POO) and weighted percent of occurrence (wPOO) (Figure 1; see Box 1 for details).

Some dietary metabarcoding studies present RRA data along with occurrence summaries, although relatively few have relied solely on information obtained from RRA (Table 1). In almost all of these studies, the number of sequences obtained per sample are converted to percentages (Figure 1a), because the absolute counts are dependent on several factors unrelated to the overall importance of the sample (amount of starting material used, DNA extraction efficiency, standardization of samples before HTS, etc.). To provide an overall diet summary, sample-specific RRA values can be averaged across samples; when doing so, each sample is given equal weight (Box 1; Figure 1b). The RRA of taxa in each sample will vary depending on genetic marker, laboratory protocol, and bioinformatic filtering strategy (Alberdi *et al.* 2017; Deagle *et al.* 2013). Ensuring laboratory methods are robust (i.e. focussing on samples with sufficient target DNA and checking replicates) and using a standardised bioinformatics pipeline without excessive filtering can help ensure RRA data are reproducible and precise (Alberdi *et al.* 2017; Deagle *et al.* 2013; McInnes *et al.* 2017a; Murray *et al.* 2015).

### Box 1 Some metrics used to summarise sequence data in dietary metabarcoding

#### Occurrence Data

Frequency of occurrence (FOO) is the number of samples that contain a given food item, most often expressed as a percent (%*FOO*). Percent of occurrence (*POO*) is simply %*FOO* rescaled so that the sum across all food items is 100%. Weighted percent of occurrence (*wPOO*) is similar to *POO*, but rather than giving equal weight to all occurrences, this metric weights each occurrence according to the number of food items in the sample (e.g., if a sample contains 5 food items, each will be given weight 1/5). Mathematical expressions are as follows:

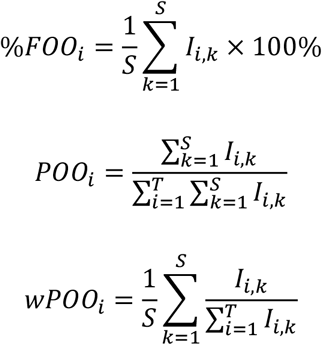

where *T* is the number of food items (taxa), *S* is the number of samples, and *I* is an indicator function such that *I_i,k_* = 1 if food item *i* is present in sample *k*, and 0 if not.

Many metabarcoding diet studies make use of both %FOO and POO (e.g. Xiong *et al*. 2017). POO provides a convenient view since each food taxon contributes a percentage of total diet (unlike %FOO which does not sum to 100%). In POO summaries samples with a high number of food taxa have a stronger influence, whereas in wPOO each sample is weighted equally (i.e. lower weighting to food taxa in a mixed meal) and this may be more biologically realistic (wPOO is the same as split-sample frequency of occurence; see Tollit *et al*. 2017 and references within).

#### Read Abundance Data

Using the sequence counts, relative read abundance (*RRA_i_*) for food item *i* is calculated as:

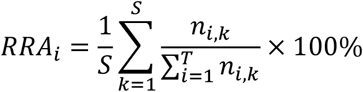

where *n_i,k_* is the number of sequences of food item *i* in sample *k*.

## 3. Does converting read counts to occurrence data solve our problems?

It is often assumed that because conversion to occurrence data moderates the impact of taxa-specific bias in marker signal, it provides a trustworthy, or at least conservative, view of diet. While it is true that occurrence-based summaries of diet are less affected by recovery bias, it is not necessarily the case that they provide a more accurate representation of overall diet. Our simulations suggest POO summaries are highly consistent but generally less accurate representation of overall diet compared to RRA summaries even when moderate taxa-specific recovery biases are present (see Box 2 for details).

### Box 2 Simulations evaluating approaches for summarising population-level diet composition

To compare how effectively occurrence and RRA methods reconstruct population-level diet we simulated HTS read counts for samples derived from a population with a fixed diet and investigated the impact of taxa-specific sequence recovery biases (Figure 2). Our simulation results are for a population with 40 food taxa in its diet, occurring in exponentially declining abundance. Sequencing was simulated for 100 scat samples assuming a mean of either 3 or 20 food taxa per sample, and assuming different sequence recovery bias scenarios: no bias, low bias or high bias. The biases introduce positive or negative biases of up to 4x and 20x (low and high biases respectively) relative to a standard. In high bias scenario a 50:50 mixture could lead to 400 fold recovery bias) Diet summaries were made using: (1) RRA; (2) POO with a 1% minimum sequence threshold. For further details see Supplementary Material (S3).

**Figure 2:**
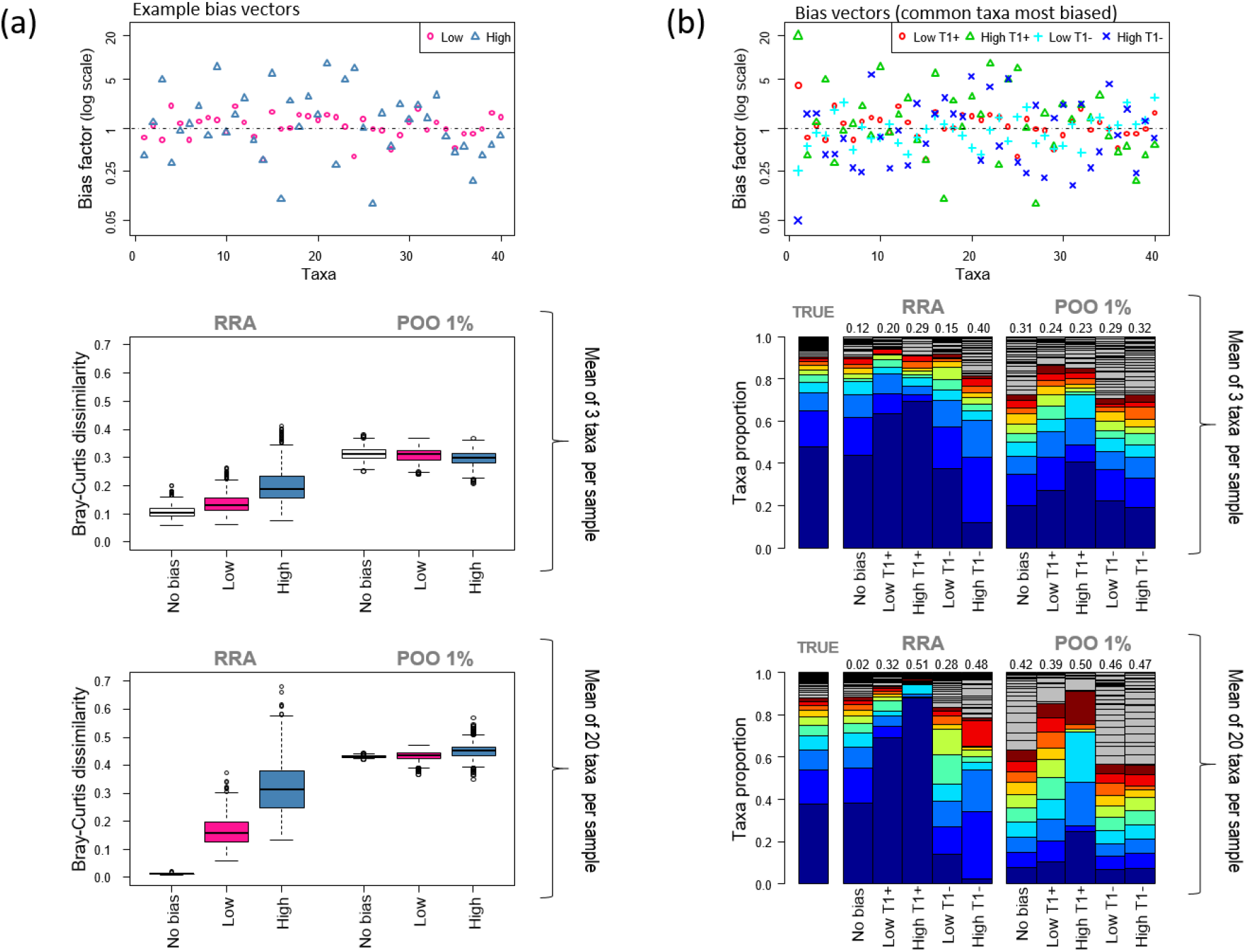
Simulation results: (a) difference between true diet proportions and estimated population diet (compared using Bray-Curtis dissimilarity metric) for RRA and POO summary methods under different bias scenarios. The first plot shows an example bias vector (for both low and high bias) used in one simulation with differential recovery values for each food taxa. The boxplots summarise results from 1000 simulations for each bias scenario where the average number of taxa per sample was 3 or 20. (b) In these simulations the most common taxa (T1) was forced to have the greatest positive bias or the greatest negative bias (low bias scenario = Low T1+ or Low T1-; high bias scenario = High T1+ and High T1-). Plots show the bias vectors and the corresponding population diet summaries are illustrated as bar plots. Numbers on top of bars are Bray-Curtis dissimilarity compared to true diet. Again, the average number of taxa per sample was 3 or 20. See Box 2 text and Supplementary Materials (S3) for details.

Overall results show that with these parameters RRA summaries were on average more accurate but had higher variance than POO summaries. POO produced more consistent estimates less impacted by recovery biases, but only outperformed RRA when the largest recovery biases corresponded to the most common food items. Both methods were more accurate when the number of food taxa per sample was small: with a small number of food taxa per sample POO estimates provide more realistic enumeration of rare items and RRA estimates are less impacted by sequence recovery biases (since biases are only expressed in the context of other taxa in a sample).

The primary drawback of occurrence datasets is that the importance of rare food taxa are often artificially inflated at the expense of food taxa eaten in large amounts, effectively flattening the rank-abundance species curves typically seen in dietary datasets (Figure 1; Box 2). This effect can be illustrated in metabarcoding data from a population-level diet study of killer whales (Figure 3). This study concluded that the whale population’s diet consisted primarily of Chinook salmon (∼80%) based on high RRA of this species in most samples (Ford *et al.* 2016). If we consider the killer whales’ diet as occurrence (POO; each food species occurrence given equal value), the view changes considerably because other salmon species and halibut frequently detected at low levels become important prey. The threshold level used to count an occurrence also impacts the relative importance of these fish prey; a lower threshold increases the importance of rare diet items (Figure 3). These different diet estimates have substantial implications when diet percentages are combined with bioenergetics estimates and consumer population size to derive estimates of prey consumption (Chasco *et al.* 2017). Another implication of rare-item inflation occurs in studies of niche partitioning. Here, the conclusion that species feed on separate resources may be inaccurate because separation may be driven primarily by partitioning of rare diet items, which are given similar weight as shared important food. In contrast, the conclusion that species overlap in their dietary niche is potentially less likely (i.e. requiring overlap in both primary and rare food items), but may therefore be more biologically meaningful when found (Clare 2014).

**Figure 3:**
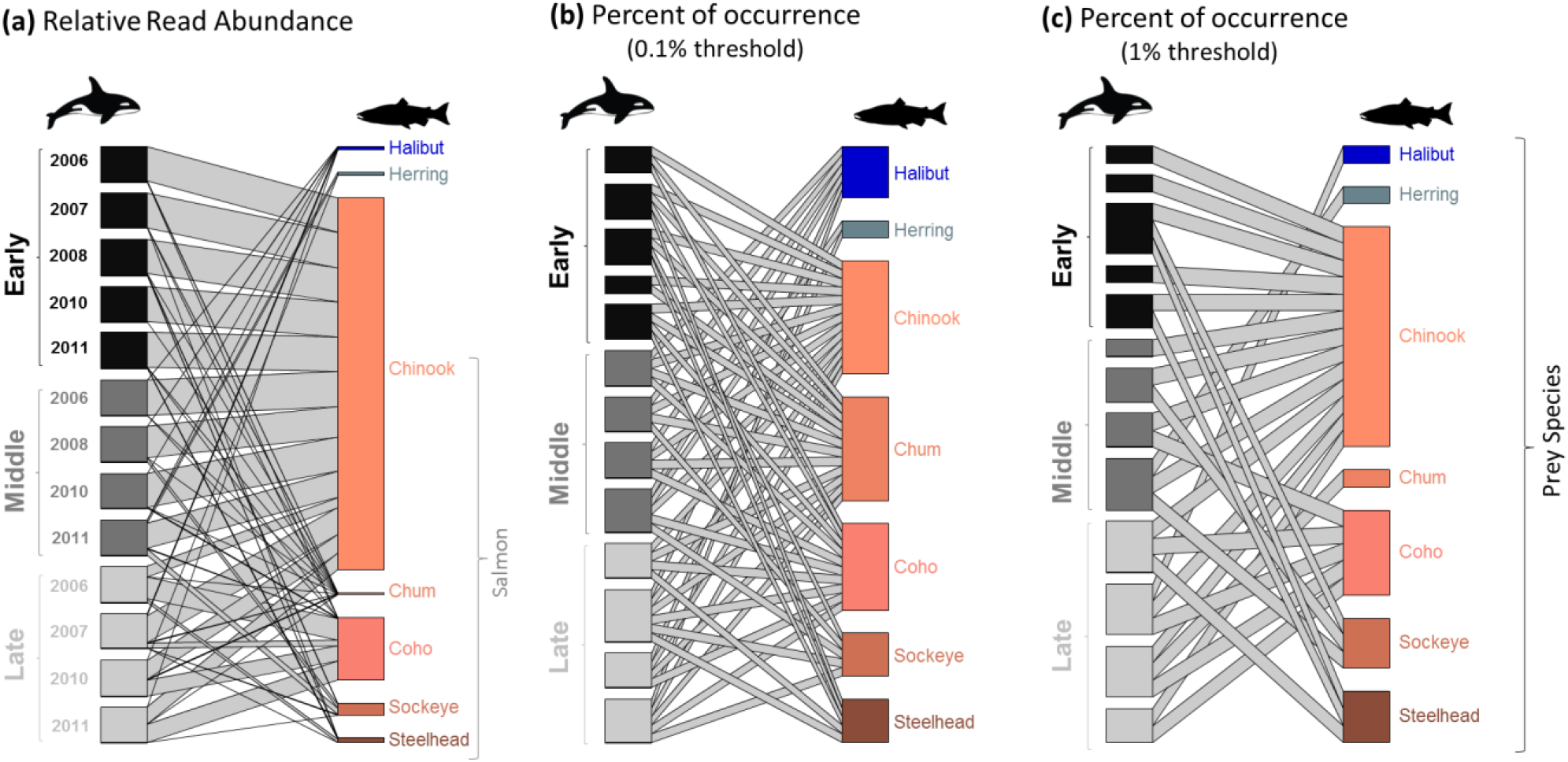
Killer whale diet in the Salish Sea illustrated with bipartite graphs constructed from data in Ford *et al.* (2016) using either (a) RRA (b) POO with a 0.1% threshold or (c) POO with a 1% threshold. Samples (DNA from faecal material) are shown on left of each plot and were pooled according to collection dates (Early, Middle, Late) in different years. The overall diet calculated by the different methods is shown on the right of each plot (includes the seven prey taxa with >1% of sequences in at least one sample). Line thickness shows contribution of taxa in each sample to the overall diet.

How much influence rare diet taxa have in overall diet estimates depends to some extent on the foraging strategy of the focal species and food distribution. In cases where small amounts of rare diet items are consumed in most feeding bouts, the importance of these items could be strongly over-estimated in occurrence-based summaries (as seen in the simulations with a high number of taxa per scat sample; Box 2). This may be the situation for some large grazing herbivores that forage continuously across a grassland, often eating relatively rare plant taxa in proportion to their availability (i.e., non-selective feeding). In contrast, when rare diet items are eaten sporadically, their DNA would be detected only occasionally and diet estimates would be more realistic. For instance, some carnivores feed sporadically, individualistically, and in discrete foraging events such that prey occurrences may provide a more meaningful indication of how often each taxon is predated (Codron *et al.* 2016). The feeding ecology of a species is reflected to some extent in the number of food taxa in individual faecal samples and this varies widely between studies (Table 1). This value provides insight into the potential impact of rare-item inflation bias. For example, in Figure 1, the zebra faecal samples have many food taxa per sample and when summarised as occurrences, these have a predictably flat rank-abundance curve; this curve would be generated regardless of the true amount of each plant consumed in each meal (Box 2).

Summaries based on occurrences become less accurate when samples are pooled (i.e. when sequence reads from individual scats are not identifiable; Clare *et al.* 2014; Deagle *et al.* 2009; Ford *et al.* 2016) because rare diet taxa present in any one of the pooled samples are weighted equally to taxa found in all of the pooled samples. The time period over which food consumption is integrated in a faecal DNA sample (influenced by gut passage time) can affect these data in a similar way, since longer integration will mean rare taxa have a greater likelihood of being present in each sample.

The inflated importance of rare sequences in occurrence summaries could also magnify some problems encountered in diet metabarcoding. There are occasions when exogenous DNA can contaminate a sample of interest. This includes field-based contamination from non-food eDNA (McInnes *et al.* 2017a), laboratory contamination (De Barba *et al.* 2014), and misassignment of sequence-to-sample during HTS (i.e. tag-jumping; Schnell *et al.* 2015). These problems will generally have less influence in RRA summaries since the real food items should dominate unless samples are very poor quality. A similar issue is the detection of secondary predation (i.e. DNA from gut contents of ingested prey). Depending on the study system and research question, secondary predation may or may not be a serious problem. However, occurrence-based datasets are expected to over-emphasise these detections and ruling out secondary predation in occurrence summaries may require information of RRA, examination of prey co-occurrence, or expert knowledge (Bowser *et al.* 2013; Hardy *et al.* 2017; McInnes *et al.* 2017b).

## 4. Does RRA actually reflect food biomass?

The relationship between proportions of biological material in a sample and sequence reads recovered by HTS has been studied in many experiments by sequencing artificial mixtures with known composition. These ‘mock communities’ are most relevant to dietary metabarcoding studies when made from food tissues similar to what is being consumed. Both mitochondrial and chloroplast DNA markers are present in multiple copies in each cell and copy number varies between tissue types (e.g. leaves versus roots; Ma & Li 2015) and physiological state (e.g. juvenile vs. gravid adult; Veltri *et al.* 1990). Getting a thoroughly homogeneous mix of tissues in a small volume suitable for DNA extractions is challenging; therefore, mixtures made from DNA extracted separately for each taxa are sometimes used (e.g. Ford *et al.* 2016; Krehenwinkel *et al.* 2017; Piñol *et al.* 2015). However, mixing purified genomic DNA will miss differences in cell density, and differences in genome size will confound results, making interpretation difficult (Piñol *et al.* 2015). Mixtures of PCR products can identify technical biases (e.g. assessing PCR primers), but miss underlying biological differences.

Conclusions from analyses of mock communities vary from no relationship to good correlations between the composition of the mixture and sequence reads (Edgar 2017; Kimmerling *et al.* 2018; Pornon *et al.* 2016). One reason for these different conclusions is that the range of concentrations analysed varies considerably across studies, from equal mixtures of a few taxa, to mixtures containing many taxa in very different abundances. A positive relationship between RRA and sample composition across a broad range of concentrations (often plotted on a log-log scale (e.g. Elbrecht & Leese 2015; Nichols *et al.* 2016)) might be missed over a smaller range. High variability between studies is also due to biotic differences in target organisms and technical differences (e.g. different barcode markers, PCR primers, sequencing platforms, etc.). This variation makes it difficult to generalise, and considerable work is required to understand the reliability of RRA in any system. Two taxonomic prey groups that have been the focus of several dietary metabarcoding studies, and for which mock communities have been examined, are fish and insects. These groups provide some insight into the expected scale of biases.

In metabarcoding of fish mixtures, conserved PCR primers are generally employed and documented recovery biases are moderate. In their killer whale study, Ford *et al.* (2016) analysed known percentages of DNA extracted from four fish species and the RRA of each fish corresponded well to input (generally within 5% of expected values) providing confidence in their conclusions. Using prey species of harbour seals Thomas *et al.* (2016) carried out a detailed study on sequence recovery from blended tissue mixtures. Various taxa (primarily fish; n=18) were sequenced in 50:50 tissue mixes with a control fish, and the extent of deviations from the control fish measured. The recovered sequences varied from 20% to 60%, a 3-fold variation in marker recovery relative to the control. A recent study looking at recovery of barcode markers from bulk samples of larval fish avoided marker amplification by directly sequencing all DNA, then bioinformatically recovering relevant marker sequences (Kimmerling *et al.* 2018). They found strong correspondence between biomass in known mixtures and sequence counts, suggesting that without PCR amplification biases, biological differences in mtDNA density between these fish are small. Even studies looking at fish environmental DNA samples have found a relationship between fish density and recovered sequence counts (Lacoursière-Roussel *et al.* 2016; Port *et al.* 2015; Thomsen *et al.* 2016).

Many studies have sequenced DNA from insect mock communities; however, rather than considering if read counts are proxies for input biomass, the focus of these studies has generally been to test if taxa can be detected (Alberdi *et al.* 2017; Clarke *et al.* 2014; Elbrecht & Leese 2015; Yu *et al.* 2012). The reason for this focus is that insect communities tend to be complex, with many rare taxa, and the recovery biases large. In studies by Yu *et al.* (2012) and Clarke *et al.* (2014), a paltry 43-76% of species known to be present in mock communities were recovered. A study that included a mixture containing equal amounts of purified DNA from 12 arthropod species (10 insects, 2 spiders), reported RRA values for half of the species were that were more than 100 times lower than expected (i.e. expected 8% and recovered at <0.08% (Piñol *et al.* 2015)). Another arthropod study found consistent relationships between percentages of DNA and RRA; however, the slope of the correlation deviated from the expected value of 1 in different insect orders and with different DNA markers, which was attributed to copy number variation (Krehenwinkel *et al.* 2017). Even a change in PCR primers used to amplify a marker from the same gene can produce very different results (Alberdi *et al.* 2017). Because of the generally poor correlation between biomass and read counts most diet studies looking at insectivorous predators focus on occurrence data (Table 1), but methodological improvements may change this (Jusino *et al.* 2017).

Diet studies incorporate more complexity than analysis of mock communities due to potential differential digestion of food taxa. Relatively few captive feeding experiments have examined how well dietary DNA counts reflect known diet, but studies have been carried out on herbivores (sheep, deer) and marine predators (penguins, seals). These have focussed on simple diets (∼2-6 diet items) and results generally show that comparisons between major and minor diet components are reflected in RRA. For example, the diet of sheep fed two plants in ratios of 0:100, 25:75, 50:50, 75:25, 100:0 had a good correlation with the percentages of DNA marker sequences amplified from rumen content (Willerslev *et al.* 2014). In a study on captive deer, >90% of the diet was made up of three plant species with two other species fed in low amounts. In this case >90% of sequences came from the three dominant taxa, but considering just these taxa, the correlation between what went in and what came out was poor (Nakahara *et al.* 2015). Similarly, in faecal samples from captive penguins fed pilchards as the majority of their diet, sequence reads from pilchards were most common in the data; however, the three other fish species fed in mass ratios 45:35:20 produced sequences counts of 60:6:34 (Deagle *et al.* 2010).

Detailed captive feeding studies examining quantitative prey DNA recovery have been carried out on captive seals and sea lions (Bowles *et al.* 2011; Deagle & Tollit 2007; Thomas *et al.* 2014). Early studies used quantitative PCR rather than HTS and found the amount of marker DNA recovered provided a reasonable indication of biomass ingested (Bowles *et al.* 2011; Deagle & Tollit 2007). A trial with harbour seals by Thomas *et al.* (2014) compared HTS data from food tissue (affected by biological and technical biases) with faecal DNA (affected by digestion as well). The scale of bias introduced by digestion was generally smaller than biases observed in undigested fish tissue mix. Since digestion bias may be in the same or opposite direction to tissue biases, the overall effect is expected to increase variance in prey-specific recovery biases compared to tissue mixes. These seal studies all excluded prey hard parts from DNA extractions, but in other systems where this may not be feasible, digestion biases could be larger. For example, faeces from insectivorous animals often contain relatively undigested hard body parts (i.e. exoskeleton). The impact on DNA recovery is difficult to assess: hard fragments will contain undigested DNA, but the DNA may not be extracted as efficiently as DNA present from soft bodied prey (Clare 2014).

Another approach to understanding how much of a signal is present in counts from DNA sequences is to compare results with other methods of diet analysis. In a study of large mammalian herbivores, Kartzinel *et al.* (2015) found a nearly one-to-one correlation between estimates of C_4_ grass (family Poaceae) consumption based on stable isotopes analyses and RRA based on metabarcoding of the chloroplast marker (trnL-P6). The use of alternative proxies for diet composition can also reveal complexities. Craine *et al.* (2015) used similar protocols to Kartzinel *et al.* (2015) but found C_4_ grass RRA to be underrepresented compared to measures based on stable isotopes. They suggested that chloroplast density scales with foliar nitrogen concentrations so that RRA values could reflect dietary sources of protein, and thus may deviate from dietary sources of biomass as represented by carbon stable isotopes. If RRA values based on this marker occasionally reflect an animal’s source of protein more closely than its source of carbon (i.e., biomass), this knowledge can enable count data to still be interpreted appropriately.

Several studies have used traditional morphological analysis of food remains to help cross-validate RRA data (Soininen *et al.* 2009; Thomas *et al.* 2017). Thomas *et al.* (2017) analysed DNA and prey hard parts in >1000 seal faecal samples, and while there were minor differences between methods in prey recovery and taxonomic resolution, both methods provided a highly similar picture of population-level diet (Thomas *et al.* 2017; Table S2). Cross-validation has the problem that all methods of diet determination are biased, so if there is disagreement the correct answer may be unclear (Soininen *et al.* 2009). However, congruence between datasets is reassuring and known biases can be taken into account when making conclusions (e.g. jellyfish are digested quickly, so occurrence in faecal DNA but not stomach contents is credible; Jarman *et al.* 2013; McInnes *et al.* 2017b). Large differences in results between methods warrant further investigation; multiple lines of independent evidence provide the strongest support for any conclusion.

Overall, assessing recovery bias between food taxa is complex, specific to a study system, and can require significant effort. In some cases, broad correlations are likely, but this cannot be taken for granted and very large biases may occur (e.g. Pawluczyk *et al.* 2015).

## 5. A view of the way forward in interpreting sequence counts

What should be considered best practice given the potential biases we have outlined in diet metabarcoding studies? First of all, we should take a step back and remember that getting estimates of the true diet of any species using any method is a challenging proposition - all methods of diet analysis have biases. A well-designed metabarcoding diet study may provide as accurate an estimate as any other approach, while also providing high taxonomic resolution, the opportunity to detect rare foods and the potential to solve otherwise intractable problems (e.g. liquid feeding). We should also remember that other classic experimental design issues, such as collecting appropriate sample sizes and getting representative samples, will potentially have a bigger impact on study outcomes than the diet estimation method. Furthermore, dietary metabarcoding has a huge variety of applications, many of which do not require highly accurate dietary proportions.

Still, we will inevitably come to a point in dietary metabarcoding studies where we need to decide how to interpret sequence counts. It is often the case that the overarching views of population-level diet are consistent regardless of how sequence counts are summarised (i.e. when commonly occurring food items are also represented by the highest number of sequences). This is most likely to be the case when faecal samples contain a limited number of food taxa (in the extreme case where there is only one taxon per sample, occurrence and RRA estimates are identical and recovery biases have no impact). However, some outcomes will depend on how we consider counts. Occurrence summaries are less affected by differential recovery of markers from food taxa, but tend to put much more weight on food consumed in small quantities and potential contaminants. RRA can potentially provide a weighting of food present in a sample based on biomass, but differential recovery of markers (especially from dominant food taxa) can impact data summaries. Our strongest recommendation is that if one approach is relied on heavily, some justification should be given for its use, and potential biases inherent in the method should be acknowledged and taken into account when drawing conclusions.

### 5.1 Using occurrence data

Many future diet studies will have almost no information on the scale of biases in the recovery of sequences from specific food taxa. The use of occurrence data may be a sensible approach, but careful consideration of the impact of this choice is still required and the bioinformatics steps taken to produce this dataset should be documented. We recommend converting counts to percentages (excluding non-food sequences from total count) and then defining a minimum sequence percentage threshold to determine occurrences. This will limit the impact of variation in read depth. The threshold is a trade-off between maximizing inclusion of real diet sequences and excluding low-level background noise (secondary predation, contamination, sequencing errors). A 1% threshold may be suitable for many situations, but when diets are extremely diverse with potentially large recovery biases (e.g. some bats species), then a much lower threshold may be justified (e.g. 0.01% in Alberdi *et al.* 2017). In these cases, ensuring contaminant sequences do not influence results requires additional vigilance (De Barba *et al.* 2014; Nguyen *et al.* 2015). Given that many of the issues we have raised regarding the use of occurrence data stem from the disproportionate influence of rarer sequences, it may seem advantageous to use a higher minimum sequence threshold (e.g. >5% constitutes occurrence). While this type of summary can provide insight, rare taxa that make up a small percentage of sequences in many samples would be missed completely (Alberdi *et al.* 2017) and taxa-specific biases in recovery also have a larger impact on these high threshold occurrence summaries (see simulations in Supplementary Material S3 comparing different threshold levels). Since the purported benefit of occurrence-based approaches is to record food taxa even when there is strong bias against them, thresholds higher than 1% cannot be generally recommended.

The sequencing depth required per sample is directly related to the minimum threshold; in diverse and/or potentially highly biased situations warranting a very low threshold (e.g. 0.01%), high numbers of reads per sample would be needed (e.g. >10000). Lower read depth is sufficient with a 1% threshold and increasing replication (biological or technical) would be preferable to having redundant sequences within samples. Even when sequence counts are not used directly, it is important these data are available as supplementary material (and ideally the sequence reads archived) with appropriate explanatory files outlining potential biases. This allows others to revisit the data and will allow insight in future comparative meta-analyses.

Summaries of data based only on occurrence information will remain appropriate in many situations. This includes dietary metabarcoding studies that use DNA from food remains in gut contents since differences in time since ingestion will have a major impact on relative number of reads recovered per taxon (Egeter *et al.* 2015; Greenstone *et al.* 2014). In studies using faecal samples, occurrence summaries will often be appropriate when food is clearly differentially digested, the sequence recovery bias is known to be high (e.g. many animals with an insectivorous diet), or this bias is unknown and results cannot be crossvalidated. Note, that this appropriateness may differ between dietary analyses of relatively similar consumers. For example, most bat diet studies only analyse occurrence data, but the bat *Myotis daubentonii* (Figure 1) has relatively low diet richness compared to other bats and RRA may be suitable (Vesterinen et al. 2016).

### 5.2 Using RRA

Incorporation of RRA into analyses will have the most benefit when individual faecal samples contain several food taxa and the same food taxa occur across many samples. In these cases, occurrence summaries may provide very inaccurate summaries (Box 2). Unfortunately RRA-based summaries from these types of samples can be most affected by recovery biases (Box 2) and careful decisions about how to interpret data are required. When there is uncertainty surrounding which method will be more accurate, presentation of results summarised with both methods is recommended. Conclusions relying heavily on RRA should include justification as to why the counts are expected to contain a roughly accurate signature. One way to justify interpretations based on RRA is through crossvalidation of the diet data with alternative methods, and this is recommended whenever possible. Alternatively, mock community experiments and/or feeding trials can be carried out, but this is feasible in a limited number of situations. In study systems where diet is relatively well known, examining biases in a small number of dominant food taxa can ensure they are not drastically over or underestimated and will lend support to using RRA information. The dominant diet items have by far the strongest impact on RRA diet summaries as significant shifts in percentages of these species will adjust percentages of all food taxa (i.e. unit sum constrained data must sum to 100%). One question that inevitably arises is at what point does “semi-quantitative” RRA information stop being useful? Our simulations indicate that even in scenarios with 20x overestimation of some food and 20x underestimation of others (i.e. in 50:50 mixtures this could lead to 400 fold recovery bias) the population-level RRA summaries often still provides a more accurate view of diet compared to POO (Figure 2). But the limits of usefulness will depend on the application. It is probable that comparisons between closely related food taxa will provide more reliable RRA data, because biological differences should be smaller and technical biases less pronounced (e.g. animal COI primer binding sites will be more conserved, or length differences in the plant trnL-P6 marker will be low). However, it is risky to make generalizations and to transfer specific methodological findings between study systems.

Further refinements to increase confidence in RRA dietary metabarcoding data are possible. Because conversion to occurrence datasets has been seen as a necessary remedy for biases in sequence recovery, there has been less incentive for researchers to test new protocols and evaluate markers on their ability to obtain accurate RRA data. While it is sensible to use standard DNA barcode markers, by ignoring information in RRA during marker development we might have inadvertently imposed limitations on the field. Fortunately, we are starting to move towards a point where markers used in different applications are better understood and alternative less-biased approaches are being explored (e.g. the use of multiple target markers (Stat et al. 2017) or PCR-free approaches (Srivathsan et al. 2016)). Inclusion of control materials in sequencing runs can also ensure consistency between experiments (Hardwick *et al.* 2017). For the most accurate diet estimates, correction factors can be developed to take into account known biological differences between taxa in mixtures (e.g. gene copy number differences; Angly et al. 2014; Vasselon et al. 2018). Such species-specific correction factors have been developed for fish, with the intent of applying them in field-collected seal diet samples (Thomas *et al.* 2016).

While the effort needed to justify the RRA approach may be challenging, the possibility of obtaining more accurate diet estimates will make it worthwhile in many situations. We have seen such effort undertaken in papers addressing broad ecological questions (Kartzinel *et al.* 2015; Willerslev *et al.* 2014), and in diet studies of marine predators, where population consumption have significant fisheries management implications (Ford *et al.* 2016; Thomas *et al.* 2017). This approach should also be possible in monitoring programs, such as those carried out on seabird diet (Jarman *et al.* 2013; Sydeman *et al.* 2017), where the long-term investment warrants the development of robust DNA-based methods that provide the best possible data.

### 5.3 Outstanding issues

There are a number of issues in the diet metabarcoding literature that have an impact on both occurrence and RRA summaries that have yet to be clearly addressed. One of these is the impact of collecting data with markers that have low taxonomic resolution (McInnes *et al.* 2017b) or collating data at higher taxonomic levels to increase certainty in taxonomic assignment (Biffi *et al.* 2017). Depending on how broad the grouping are, occurrence summaries may not be very informative as many occurrences are potentially pooled. For RRA it is unclear whether pooling counts from multiple taxa will nullify fine-scale stochasticity in recovery biases, or magnify lineage-specific biases. A related issue is how to summarise data from diet metabarcoding studies using multiple markers. When markers are targeting the same food taxa, either additive (i.e. include detections by any marker) or restrictive strategies (only include food detected by all markers) could be logically applied in occurrence and RRA summaries (Alberdi *et al.* 2017). The situation is even more complex when a “universal” primer set is used to define the broad diet and group-specific primers subsequently improve taxonomic resolution for particular groups (e.g. a marker targeting all plants together with several that offer greater resolution for specific plant families). Errors based on the universal marker will be propagated when attempting to incorporate data from the other primer sets (i.e. if the grass family is estimated to be 20% of a diet instead of the true 40%, then the perceived importance of each grass species is reduced).This problem can be avoided to some extent by reporting each component separately, but this provides an unsatisfactory synthesis for omnivorous and other species with a very diverse diet that can only be characterised with several markers (De Barba *et al.* 2014). Studies that use a marker capturing only one component of the diet need to be very clear that the results comprise an unknown amount of the total diet.

Simulations such as the ones outlined in this paper can help establish which scenarios are most sensitive to biases (from either occurrence or RRA). When informed by experimental work to assign an error range to each parameter, and combined with sensitivity analysis, this can identify which sources of bias have the largest impact on conclusions. Some of the details we have focussed on may be inconsequential for many studies and we have not considered the effect of alternate summaries on downstream applications. For example, it would be very interesting to see how switching between occurrence and RRA datasets affects outputs in the context of food web studies (Roslin & Majaneva 2016).

The ultimate test for how to deal with sequence counts in HTS diet analyses will remain in empirical studies. We hope this opinion piece will be a starting point to highlight the need to consider all sources of bias and to justify the methods used when confronting count data in metabarcoding studies. We also hope that this critique is not discouraging to researchers approaching this new and rapidly developing area of research, as no single study should be rightly expected to address all issues arising from DNA-based diet analyses. Instead, our aim is to encourage researchers to continue to addressing methodological challenges, and acknowledge unanswered questions to help spur future investigations. As the field matures, we envisage publication standards will emerge to provide the most robust diet data and provide an accurate indication of the uncertainty associated with dietary assessments.

## Data Accessibility

All data in figures is either publically accessible or will be deposited in Dryad along with R scripts to produce the figures (including simulations).

## Author Contributions

All Authors contributed ideas and to the writing of the paper

